# Structural, functional and molecular dynamics analysis of *cathepsin B* gene SNPs associated with tropical calcific pancreatitis, a rare disease of tropics

**DOI:** 10.1101/308932

**Authors:** Garima Singh, MSK Jayadev, Rinku Sharma, Basharat Bhat, CH Madhusudhan, Ashutosh Singh

**Affiliations:** School of Natural Sciences, Department of Life Sciences, Shiv Nadar University, Greater Noida, UP, India; Head, Department of Surgical Gasteroenterology, Osmania General Hospital, Hyderabad, India.

**Keywords:** Molecular dynamics, Protein simulation, *cathepsin B*, cysteine protease, tropical calcific pancreatitis, pancreatitis

## Abstract

Tropical Calcific Pancreatitis (TCP) is a neglected juvenile form of chronic non-alcoholic pancreatitis. Cathepsin B (CTSB), a lysososmal protease involved in cellular degradation process, is recently been studied as a potential candidate gene in the pathogenesis of TCP. According to *cathepsin B* hypothesis, mutated CTSB can lead to premature intracellular activation of trypsinogen, which is a key regulatory mechanism in pancreatitis. So far, CTSB mutations have been studied in pancreatitis and neurodegenerative disorders but little is known about the structural and functional effect of variants in CTSB. In this study, we investigated the effect of single nucleotide variants (SNVs) associated with TCP, using molecular dynamics and simulation algorithms. There were two non-synonymous variants in the coding region (L26V and S53G) of CTSB, located in the propeptide region. We tried to predict the effect of these variants on structure and function using multiple algorithms: SIFT, Polyphen2, Panther, SDM sever, i-Mutant2.0 suite, mCSM algorithm and Vadar. Further, using databases like miRdbSNP, PolymiRTS and miRNASNP, two SNPs in 3’UTR region were predicted to affect the miRNA binding sites. Structural mutated models of nsSNP mutants (L26V and S53G) were prepared by MODELLER v9.15 and evaluated using TM-Align, Verify 3D, ProSA and Ramachandran plot. The results showed that the models (L26V and S53G) were of high accuracy. The 3D mutated structures were simulated using GROMACS 5.0 to predict the impact of these SNPs on protein stability. The results from in silico analysis and molecular dynamics simulations suggested that these variants in the propeptide region of *cathepsin B* could lead to structural and functional changes in the protein. Hence, the structural and functional analysis results have given interim conclusions that these variants can have deleterious effect in TCP and thus should be screen in samples from all TCP patients to decipher its distribution in patient population.

## Introduction

Pancreatitis is a multifactorial, heterogeneous disease whose etiologies are still enigmatic. It is an inflammatory condition leading to morphological changes in the pancreas that causes pain and functional abnormalities. Alcohol(1), malnutrition(2), gallstones(3), familial clustering (4) and sometimes severe infections (5) have been seen to be the major cause of pancreatitis. Pancreatitis is broadly classified (6) as acute and chronic. Tropical Calcific Pancreatitis (TCP) (7) is a juvenile form of chronic calcific non-alcoholic pancreatitis. It is a form of Idiopathic Chronic Pancreatitis (ICP), mostly reported in the developing tropical countries. TCP proliferates in childhood and persist in adolescence. The phenotypic heterogeneity (8) includes abdominal pain, ductal dilation, large pancreatic calculi and pancreatic atrophy. The genetic heterogeneity is yet to be explored. Fibrocalculous pancreatic diabetes (FCPD) (9),a unique form of diabetes, is the characteristic secondary feature of TCP.

The pathophysiology of pancreas is composed of exocrine functions for digesting food and endocrine function for glucose homeostasis. Trypsinogen, cathepsins, serine proteases, calcium sensing receptors are some of the essential genes for regulation of mechanisms operating in pancreas. According to trypsin-centered theory of pancreatitis, trypsinogen, a key zymogen in pancreatic juice and a key regulator of digestion, exhibits a premature activation in acinar cells of pancreas. This lead to activation of other zymogens in the pancreas, leading to inflammation and autodigestion of pancreas, thus Pancreatitis. But, TCP, an enigmatic form pancreatitis without a cause, is increasing the mortality rate in tropical countries.

Human *cathepsin B* (catB, E.C 3.4.22.1), a lysosomal cysteine proteases is involved in several cellular processes like protein degradation, extracellular matrix degradation, regulatory mechanisms, cell death, autophagy, antigen representation (10). Cathepsin B is synthesized as an inactive proenzyme (procathepsinB) and is activated by other proteases and autocatalytic processes (11). Signal sequence and post-translational glycosylation (12) along with mannose-6-phosphate receptor mediated endocytic pathway, targets *procathepsin B* to endosomes/lysosomes(13). *Procathepsin B* has an N-terminus propeptide of 62 amino acid length from Arg-Lys. Propeptide exhibits an essential role in the processing and maturation of *cathepsin B*. It act as (a) scaffold for catalytic domain during protein folding (b) intracellular trafficking of *procathepsin B* to lysosome after N-terminal glycosylation and phosphorylation (c) high affinity reversible inhibitor for the premature activation of zymogen (14). Mutations affect different regions of *cathepsin B* protein but how these variants affect the function of *cathepsin B* is yet to be studied.

In this study, we try to comply with the “*cathepsin B* hypothesis”(15) in tropical calcific pancreatitis. According to this hypothesis, *cathepsin B* plays an essential role in the premature activation of trypsinogen in the pancreas, largely due to colocalization of *cathepsin B* and zymogens.(16) The reason behind this colocalization is yet to be deciphered which can be due to mutations in *procathepsin B* or in the molecules associated with the trafficking of *procathepsin B* in diseased state. Taking this as a background, we studied the effect of two non-synonymous mutations in the coding region (propeptide region [18-79aa]) of *procathepsin B*, screened in the TCP patients by molecular dynamics and simulation studies. Also, we predicted, using various in-silico algorithms, that variants present in 3’UTR region are associated with miRNA binding sites, and hence they are significant.

We collected the natural variants data of *cathepsin B* (17) associated with tropical calcific pancreatitis after intensive data mining, for in-silico analysis, to determine the effect of these variants on the three dimensional structure and stability of *procathepsin B* as well as on the function of protein. *Procathepsin B* gene was completely characterize and the 3D-structure of the mutants (L26V and S53G) were predicted using MODELLER 9.15 (18). The predicted structure of each mutant was further evaluated by Verify-3D (19), PROSA (20) and Ramachandran plot (21). Afterwards, predicted mutant PDB structures were aligned to the wild-type PDB structure using TM-Align (22). Plausible structural effect of mutations on the biochemical and physiochemical properties of protein was inferred using available bioinformatics tools and algorithms like SIFT (23), PolyPhen2 (24), Panther (25), ProtParam (http://web.expasy.org/protparam/) and Vadar v1.8 (26). The stability of mutated protein models was predicted using webservers like, SDM (27), I-Mutant suite (28) and mCSM (29). Thereafter, energy minimization and molecular simulations were performed for predicted mutated structures, using GROMACS (version 5.0) (30). Molecular simulations predicted the instability of mutated proteins. Further, SNVs in 3’UTR region were predicted to have miRNA binding sites, by using databases like miRdSNP (31), PolymiRTS 3.0 (32), miRNASNP (33).

Evidential results from the structural analysis and comparison of 2D and 3D mutated models of *cathepsin B* to wild-type protein has implicated the potential role of these variants in the pathobiology of tropical calcific pancreatitis. This is the first attempt to structurally and functionally characterize the variants found in human *cathepsin B* protein screened in TCP patients.

## Materials and Methods

### Data curation

The single nucleotide polymorphisms (SNPs) was mined after intensive literature survey on tropical calcific pancreatitis. The mRNA accession number, NM_147781.2, NM_001908.3 and protein accession number, NP_680092.1 of gene *cathepsin B* (CTSB) is used in our computational analysis. The SNP data was retrieved from literature (PMID16492714) (17). The workflow for the computational analysis performed to decipher the significance of SNPs is depicted in Fig 1.

**Fig 1:**
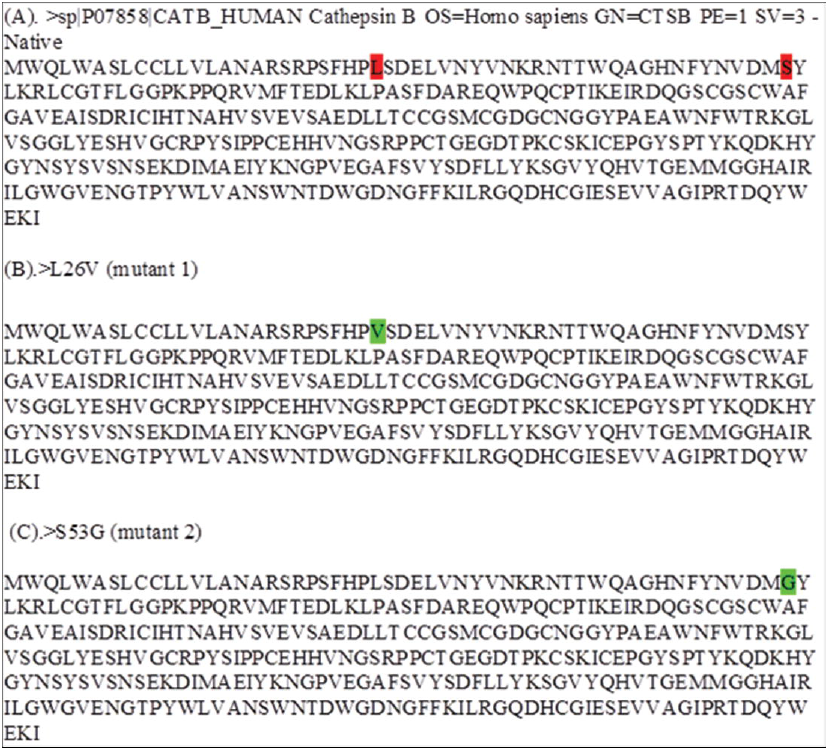
Workflow to identify the potential effect of SNPs

### Sequence retrieval and Alignment

The sequence of *cathepsin B* (CTSB) was retrieved from UniProt database-P07858 (CATB_HUMAN). The non-synonymous variants (L26V and S53G) were manually inserted in the wild-type protein sequence for further analysis.

### Homology modeling

The two ns SNPs (L26V and S53G) are modelled to analyse the structural effect of variants. The position specific iterated blast program (PSI-BLAST) with protein databank database (PDB) and default advanced settings, was used to find the template for homology modelling. MODELLER 9.15 was used to build mutated models of *procathepsin B*. The best predicted models according to lowest value of DOPE score (Discrete Optimized Protein Energy) was used for further evaluation and analysis. The predicted models as L26V and S53G were annotated and evaluated by using Verify-3D and ProSA servers. After these models pass the respective thresholds, ramachandran plot was evaluated. TM-score and root mean square deviation (RMSD) values of mutant structures (L26V and S53G), were calculated with respect to wild-type by using TM-Align.

### Non-synonymous SNP Analysis

The two dimensional structure of wild-type and mutated protein were calculated using PSIPRED v3.3. The functional effect of mutations is predicted using following algorithms: SIFT, Polyphen2 and Panther. Another tool, ProtParam was used to calculate the hydropathicity or the GRAVY (grand average of hydropathicity) score of mutated *procathepsin B*. Hydrogen bond length and the rotational angles of main chain hydrogen bonds is a significant descriptor to locally study the confirmation and dynamics. Therefore, to calculate the altered hydrogen bonding patterns in the mutated three dimensional *procathepsin B* structures, Vadar v1.8 program was used. This program calculates the H-bond distances in main chain, side chain and the bond angle between the main chain residues. Further, the stability of the mutated models was calculated by SDM server, I-Mutant suite, mCSM webservers, which are free energy (ΔΔ*G*) calculator algorithms.

### Molecular dynamics simulation

Molecular dynamic simulation was performed with Gromacs-5.0 package on the native (PDB ID: 3PBH) (35) and mutant structures (S53G and L26V). This computational investigation was done with a view point to examine if these single nucleotide variants might lead to changes in surface properties or distort the protein orientation. The protein molecule was solvated in a dodecahedron box with SPC216 water molecules at 1.5 Å marginal radiuses. The system was made neutral by adding required number of sodium (Na) and chloride (Cl) ions. Subsequently, the molecular system was subjected to steepest distance energy minimization until reaching the criterion of 1000Kj/mol with OPLS-all atom force field. Berendson temperature coupling method was used to regulate the temperature inside the box at 300k. Isotropic pressure coupling was performed using Parinello-Rahman method and the pressure of the system was maintained at 1 bar. LINCS algorithm was used to constraint bond lengths including H-bonds. Van der Waals and Coloumb interactions were truncated at 1nm and Particle Mesh Ewald method was used to compute electrostatic interaction. Finally, simulation was performed for 35ns. The structural deviations between native and mutated structures were subjected to comparative analysis by computing RMSD (Root mean square deviation) and RMSF (Root mean square fluctuation). The trajectories were analysed and finally protein compactness was studied by calculating radius of gyration.

### Analysis of SNPs in UTR region

5’UTR and 3’UTR regions in a gene play crucial role to regulate gene expression at post translational level. UTRs regulate the exit of mRNAs from nucleus, translation efficiency, subcellular localization and mRNA stability (34). The effect of SNPs in UTR regions were analysed using databases like miRdSNP, PolymiRTS and miRNASNP.

## Results

### Data curation

The SNPs associated with TCP were extracted from literature and is tabulated in Table 1. The SNPs are further categorized and is shown in Fig 2. There are total 23 SNPs in CTSB gene. The non-coding region includes 20 SNPs (1 deletion, 6 in 5’UTR, 2 in 3’UTR and 11 in introns). Coding region has 2 missense variants and 1 synonymous variation.

**Fig 2.**
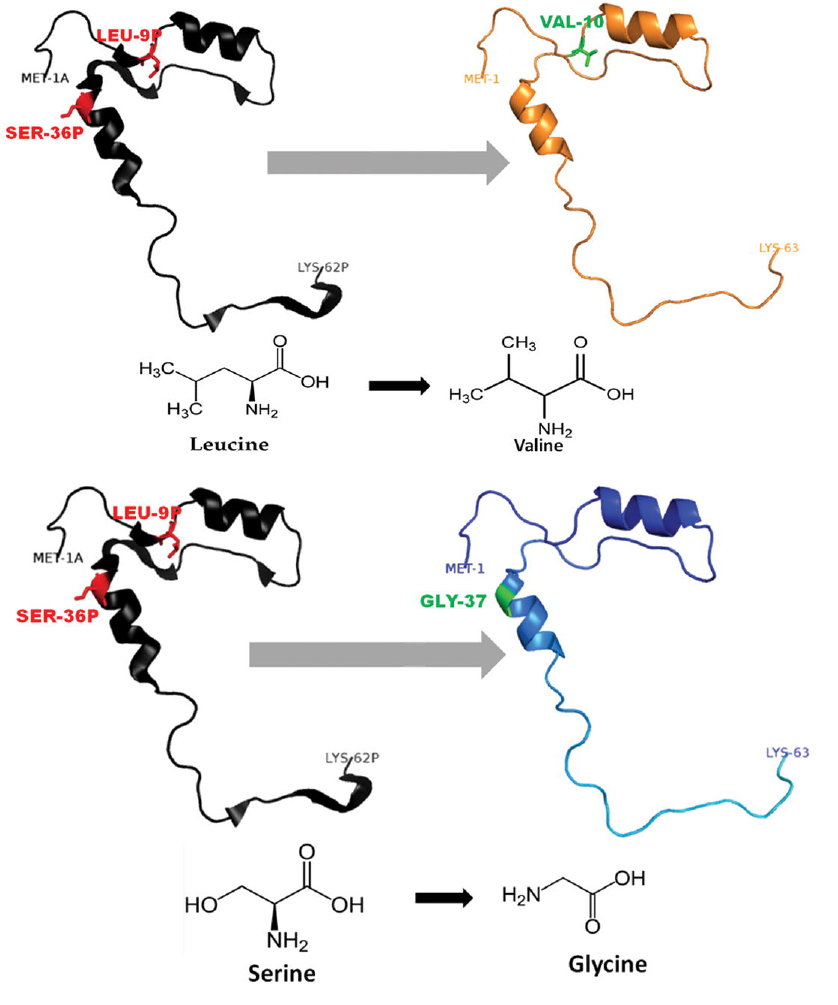
Classification of SNPs in *cathepsin B* protein retrieved from literature.

**Table 1.** The Single Nucleotide Polymorphisms in *cathepsin B* protein mined from literature (PMID: 16492714). The SNP information is with respect to Ref Seq sequence ID: NT_077531.5 and dbSNP Build 150.

| Ref rsID/ss ID | Position | Type | CDS position(relative to CDS start) | CDS Allele change | Protein Position | Residue change |
| --- | --- | --- | --- | --- | --- | --- |
| - | Exon 1(5'UTR) | Non coding | 14609 | C>A | - | - |
| - | Intron1(5'UTR) | Non coding | 14520 | G>C | - | - |
| - | Intron1(5'UTR) | Non coding | 14453 | G>A | - | - |
| rs1293311 | Intron1(5'UTR) | Non coding | 14425 | C>A | - | - |
| rs2645415 | Intron1(5'UTR) | Non coding | 11083 | T>C | - | - |
| - | Exon 2(5'UTR) | Non coding | 10927 | C>G | - | - |
| rs4292649(rs12338) | Exon 3 | Non-synonymous coding(Missense) | 76 | C>G | 26 | L>V |
| rs1293293(rs1122182) | Intron 3 | Non-coding | 335 | A>T | - | - |
| - | Intron 3 | Non-coding | 394 | G>A | - | - |
| rs1293292 | Intron 3 | Non-coding | 595 | C>T | - | - |
| rs1293291 | Intron 3 | Non-coding | 663 | T>C | - | - |
| rs1803250 | Exon 4 | Non-synonymous coding(Missense) | 790* | A>G | 53 | S>G |
| rs2272766 | Intron 5 | Non-coding | 2609 | C>T | - | - |
| rs13332 | Exon 6 | Synonymous coding | 4383* | A>C | 140 | T>T |
| - | Intron 6 | Non-coding | 4451 | G>C | - | - |
| rs1736090 | Intron 6 | Non-coding | 4735 | A>G | - | - |
| rs1692819 | Intron 7 | Non-coding | 5516 | C>T | - | - |
| rs2294139 | Intron 7 | Non-coding | 5522 | C>A | - | - |
| rs3215434 | Intron 7 | Non-coding(Deletion) | 5581-5582 |  | - | - |
| - | Intron 7 | Non-coding | 5622 | C>G | - | - |
| rs2294138 | Intron 8 | Non-coding | 5825 | G>A | - | - |
| rs709821 | Exon 11(3'UTR) | Non-coding | 8370* | C>G | - | - |
| rs8898 | Exon 11(3'UTR) | Non-coding | 8422* | A>G | - | - |

### Sequence retrieval and Alignment

The protein sequence (Table 2) was retrieved from Uniprot database and the desired variants were manually inserted in the sequence. The Uniprot ID is P07858 (CATB_HUMAN). The sequence annotations are shown in Table 3. The sequences of the wild type and the mutant are described in Fig 3.

**Fig 3.**
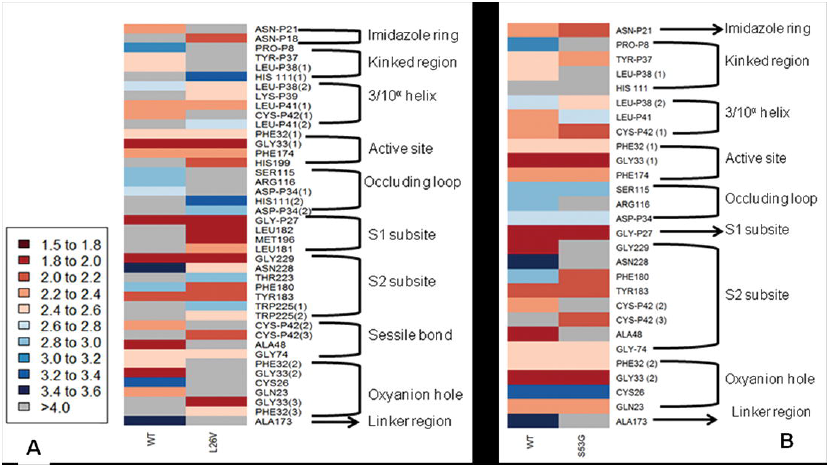
Fasta alignment of wild and mutants *procathepsin B* retrieved from Uniprot database. (A) Fasta sequence of wild-type *procathepsin B* (NP_680092.1, Isoform 1) retrieved from Uniprot database (ID: P07858). The wild type amino acids are highlighted red (B) Fasta sequence of the mutant Leu26Val (L26V) manually substituted in the wild-type protein sequence. The substituted amino acid, valine is highlighted as green. (C) Fasta sequence of mutant Ser53Gly (S53G) manually substituted in the wild-type protein sequence. The substituted amino acid, glycine is highlighted as green.

**Table 2:** Summary of non-synonymous coding variants retrieved from literature

| Sequence ID | Position/Region | Amino acid change |
| --- | --- | --- |
| NP_680092.1(NCBI Accession No.) | 26/Exon3 | Leu to Val (L26V) |
| P07858 (Uniprot ID) |  |  |
| NP_680092.1(NCBI Accession No.) | 53/Exon 3 | Ser to Gly (S53G) |
| P07858 (Uniprot ID) |  |  |

**Table 3:** Summary of annotation of structures of proteins

| Sequence | Annotation |
| --- | --- |
| Wild Type | >WT (Fig 3A) |
| Mutant 1 | >L26V (Fig 3B) |
| Mutant 2 | >S53G (Fig 3C) |

### Homology modelling

The 3D-structure of mutants (L26V and S53G) proteins was predicted after template searching by PSI-BLAST. The protein sequence (NP_680092.1) of *procathepsin B* was used as query and the resulting templates were then filtered. Finally, X-ray crystal structure of human *procathepsin B* (PDB ID: 3PBH) (35) with 2.5 Å resolution was used as a template for homology modelling. The DOPE scores, TM-scores and RMSD of the predicted best models by Modeller 9.15 are shown in Table 4. The stereochemical properties of the mutated *procathepsin B* structures were evaluated using Ramachandran plot from RAMPAGE. The plot defines the amino acids in favoured, allowed and outlier regions in the mutated structures as well as in the wild-type *cathepsin B* structures in Table 5. Structure validation of the predicted models was done by Verify-3D and it was observed that 99.05% amino acids have average 3D-1D protein score in a 21 residue sliding window >=0.2 in L26V mutated model and 97.16% amino acids have average 3D-1D protein score in a 21 residue sliding window >=0.2 in S53G mutated model. Additionally ProSA web server was used to evaluate the quality of predicted mutated models. The Z-score of L26V model is −7.32 and S53G model was −7.47 which are within the acceptable range of X-ray and NMR studies. The interaction energy analysed by ProSA tool is negative for maximum residues in L26V and S53G predicted models, in a sliding window of 10 and 40 respectively. Since the mutations are present in the propeptide region of *procathepsin B*, the mutated propeptide structures are shown in Fig 4 and Fig 5.

**Fig 4:**
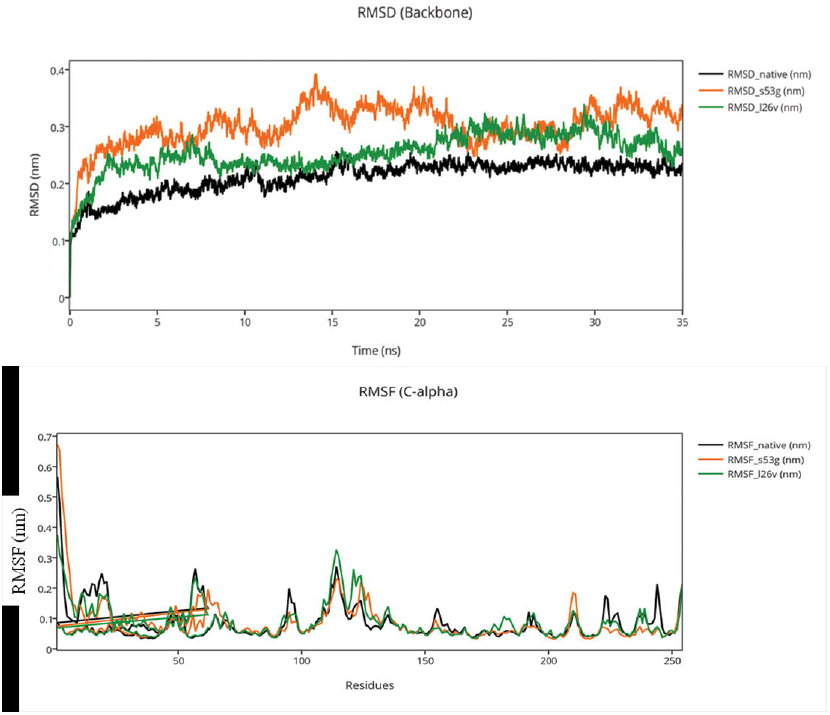
Mutated (orange color) and wild-type (black color) propeptide models are shown. The mutation, L26V is shown in sticks, which is equivalent to LEU9P-VAL10 in PDB files. Visualization and numbering is done using PyMOL tool. Note: Numbering of amino acids in wild type PDB (3PBH) file differ to that of mutated models because propeptide and peptide regions are numbered separately in the published wild-type PDB file.

**Fig 5:**
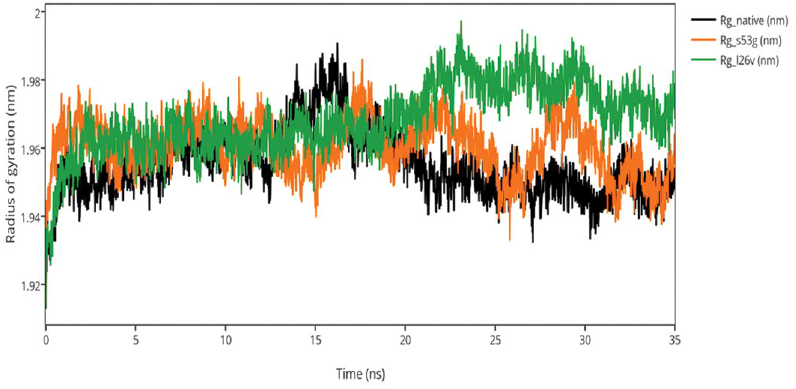
Mutated (Blue color) and wild-type (black color) propeptide models are shown. The mutation, S53G is shown insticks as SER36-GLY37 in PDB files. Visualization and numbering is done using PyMOL tool. Note: Numbering of amino acids in wild type PDB (3PBH) file differ to that of mutated models because propeptide and peptide regions are numbered separately in the published wild-type PDB file.

**Table 4:** DOPE scores after homology modelling by Modeller 9.15 of mutants (L26V and S53G) and the structure alignment scores (TM-score and RMSD) of the CTSB mutant models with wild-type, 3PBH structure.

| Predicted<br>mutant model | DOPE Score | TM-score | RMSD |
| --- | --- | --- | --- |
|  | Modeller 9.15 | TM-Align |  |
| L26V | -34105.02344 | 0.99941 | 0.16 |
| S53G | -34285-52344 | 0.99973 | 0.17 |

**Table 5:** The Ramachandran plot analysis of mutated models.

| Models | Favoured<br>region | Allowed<br>region | Outlier<br>region |
| --- | --- | --- | --- |
| L26V | 92.7% | 5.1% | 2.2% |
| S53G | 91.7% | 6.0% | 2.0% |

### Non-Synonymous SNP Analysis

The functional effect of the mutations predicted by using algorithms described above are tabulated in Table 6. The S53G mutation was observed to have deleterious effect on the function of *procathepsin B* while average analysis of L26V mutation implicit that the mutation is insignificant. The GRAVY scores of wild-type WT is −0.470 and the mutants (L26V and S53G) mutants is −0.469 for both. Thus, there is no significant effect of mutations on hydropathicity of the protein. The comparative analysis of hydrogen bond lengths and the rotational angles between WT and mutants were calculated by using Vadar v1.5 server, at 10 different regions of protein, playing imperative role in the functioning of *procathepsin B*. It was observed remarkable difference in the H-bond lengths and Bond angles between WT and mutants (S53G and L26V) as displayed in Fig 6. Hence, it can be interpreted that mutated structures alters the binding between residues, which can further prove to be deleterious in downstream signalling of *procathepsin B*. The data is shown in Fig 5. The free energy (ΔΔ*G*) of the mutated structures calculated by SDM server, I-Mutant 2.0, and mCSM webservers indicates destabilization of mutated proteins (Table 7).

**Fig 6:**
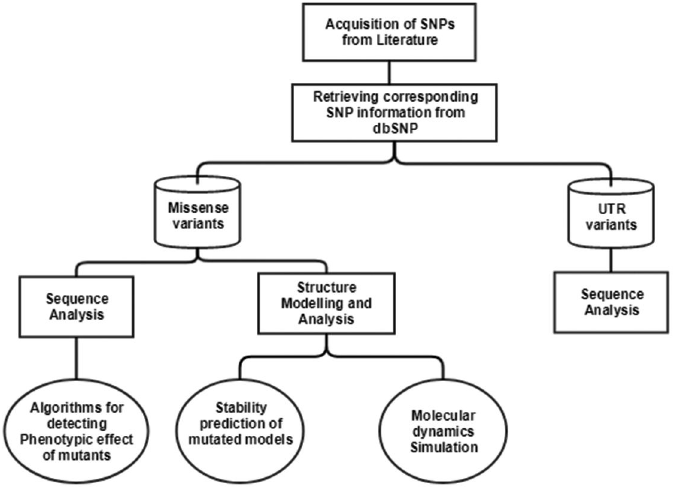
The comparative analysis of H-bond length between wild-type (WT) *procathepsin B* protein (PDB ID: 3PBH) and **A**. mutated structure (L26V) having a mutation at 26^th^ amino acid (Leucine to Valine) in propeptide region **B**. mutated structure (S53G) having a mutation at 53^rd^ position (Serine to Glycine) in propeptide region. The colour key ranges from 1.5Å to 3.5Å with red as strong H-bonding and blue as weak H-bonding. Wide spaces indicate the absence of H-bonds at that position.

**Table 6.** Prediction of functional effect of mutations by using different algorithms.

|  |  |  | SIFT |  | Polyphen 2 |  | Panther |  |
| --- | --- | --- | --- | --- | --- | --- | --- | --- |
| rsID | Allele change | AA change | Score | Prediction | Score | Prediction | Score | Prediction |
| rs4292649(rs12338) | C>G | L26V | 0 | Affect protein function | 0.01 | Benign | 1629 | Probably damaging |
| rs1803250 | A>G | S53G | 0.02 | Affect protein function | 0.06 | Benign | 750 | Probably damaging |

**Table 7.** Prediction of protein (*procathepsin B*) stability upon mutation. The calculated free energy (ΔΔ*G*) changes is in kcal/mol. ΔΔ*G* >0 indicates stabilization while ΔΔ*G*<0 indicates destabilization. The mutations are destabilizing according to these prediction algorithms.

| Algorithm | S53G | L26V | Effect |
| --- | --- | --- | --- |
| SDM | -1.174 | -0.94 | Destabilizing |
| I-Mutant 2.0 | -1.48 | -1.93 | Destabilizing |
| mCSM | -1.063 | -1.638 | Destabilizing |

### Analysis of SNPs in UTR region

UTRs play an important role in mRNA processing during post-translational mechanism. Hence the SNPs in the UTR region can significantly affect the functionality of UTRs. The 3’UTR region is essential for microRNA (miRNA) binding which can lead to degradation or transcriptional suppression of mRNA and thus further affect the downstream processing. The databases miRdbSNP, PolymiRTS, miRNASNP, were used to predict the significance of SNPs in 3’UTR region (Table 8). The 2 SNPs present in 3’UTR region, rs709821 and rs8898 were predicted to be present in miRNA binding sites and therefore are significant. The SNP rs8898 was predicted to be loss of functional mutation by PolymiRTS database and hence it is significant and can be deleterious.

**Table 8:** The SNPs in 3’UTR region of CTSB protein were predicted to have functional significance since they are associated with miRNA target sites.

| rsID | Region | Allele change | miRdSNP | PolymiRTS | miRNASNP |
| --- | --- | --- | --- | --- | --- |
| rs709821 | UTR-3 | C>G | hsa-miR-186 | - | - |
|  |  |  | hsa-miR-339-5p | - | - |
|  |  |  | hsa-miR-7 | - | - |
|  |  |  | hsa-miR-214 | - | - |
|  |  |  | hsa-miR-431 | - | - |
|  |  |  | hsa-miR-186 | - | - |
|  |  |  | hsa-miR-320a | - | - |
|  |  |  | hsa-miR-320d | - | - |
|  |  |  | hsa-miR-320c | - | - |
|  |  |  | hsa-miR-320b | - | - |
|  |  |  | hsa-miR-96 | - | - |
|  |  |  | hsa-miR-1271 | - | - |
| rs8898 | UTR-3 | A>G | hsa-miR-339-5p | hsa-miR-10a-5p | hsa-miR-10a-5p |
|  |  |  | hsa-miR-186 | hsa-miR-10b-5p | hsa-miR-10b-5p |
|  |  |  | hsa-miR-7 | hsa-miR-339-5p | hsa-miR-339-5p |
|  |  |  | hsa-miR-214 | hsa-miR-4421 |  |
|  |  |  | hsa-miR-431 | hsa-miR-5699-3p |  |
|  |  |  | hsa-miR-186 | hsa-miR-6747-3p |  |
|  |  |  | hsa-miR-320a | hsa-miR-6752-3p |  |
|  |  |  | hsa-miR-320d |  |  |
|  |  |  | hsa-miR-320c |  |  |
|  |  |  | hsa-miR-320b |  |  |
|  |  |  | hsa-miR-96 |  |  |
|  |  |  | hsa-miR-1271 |  |  |

### Molecular dynamics simulations

The comparative analysis of trajectories by calculating RMSD, RMSF and Radius of gyration after MD simulation of 35ns for both native and mutants was performed. Interestingly, it was observed from RMSD of backbone residues that both mutated structures (S53G and L26V) were distinctly deviated from the native structure (PDB ID: 3PBH). To infer whether the mutations affected the dynamic behaviour of each residue or not, RMSF of Cα-atoms was calculated. There was fluctuation observed in mutated protein structures as compared to native structure. The protein compactness was determined by radius of gyration (Rg). It was observed that Rg of mutant structures are distinctly fluctuated as compared to native structure all throughout the simulation.

**Fig 7.**
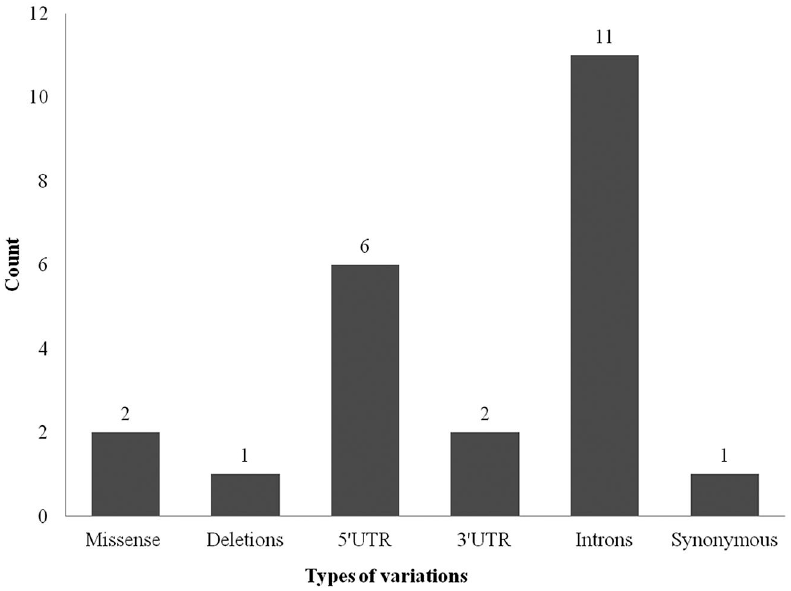
MD Simulation. A. Comparative RMSD analysis shows that the mutated structures (s53g and l26v) are distinctly deviated from native structure after simulating the proteins for 35ns. B. RMSF of C-α residues were not fluctuated much, all throughout the simulation of mutated protein structures (s53g and l26v). C. Comparative analysis of Radius of gyration (Rg) of mutated (s53g and l26v) and native proteins depicts that protein compactness is varied in mutated proteins.

## Discussion

Tropical Calcific Pancreatitis (TCP) has a distinct morphological characteristics with undefined etiology. *Procathepsin B* and activated glycosylated form of *cathepsin B* is present in acinar cells of pancreas (36). The propeptide region of *procathepsin B* (Arg1-Lys 62) i.e the N-terminal part inhibit the activity of *cathepsin B* in the pancreas, thereby regulating its premature activation, also act as a scaffold for protein folding and as chaperone for endosome/lysosomal trafficking. L26V and S53G are the two missense variants observed in the propeptide region of *cathepsin B* protein in TCP patients. The *in silico* SNP analysis of the mutated protein sequence resulted in alteration of secondary structure, thereby predicting an inimical folding effect on the protein. The phenotypic effect of the mutations was calculated using sequence analysis algorithms and it was observed that at least one algorithm indicated a deleterious effect of these mutations on protein functioning. The free energy (ΔΔ*G*) calculations of mutated proteins by varied algorithms indicated that mutations are destabilizing. Further, the comparative analysis of H-bond distances between mutated and native 3D-structure of *procathepsin B* provided a unique information about the structural characteristics of motifs around main chain H-bonds which are altered in mutant protein, thereby affecting the function of protein (37). Additionally, MD simulation of mutated and native protein indicated that the mutations distinctly deviate the structural confirmation of *procathepsin B*, which can have deleterious effects in downstream signalling mechanism. Thus, the structural and functional analysis of mutated *procathepsin B* predicts the significance of mutations in propeptide region of *cathepsin B*. These results will provide a lead towards designing the experimental research on the mutations involved in pathogenesis of TCP to understand the disease pathogenesis.

## Acknowledgements

AS is thankful to Shiv Nadar University for providing all the necessary support to perform this study.

## References

1. Pandol SJ, Gorelick FS, Gerloff A, Lugea A. Alcohol abuse, endoplasmic reticulum stress and pancreatitis. Dig Dis. 2010; 28:776–82.

2. Witt H, Bhatia E. Genetic aspects of tropical calcific pancreatitis. Rev Endocr Metab Disord. 2008; 9(3):213–26.

3. Levy P, Dominguez-Munoz E, Imrie C, Lohr M, Maisonneuve P. Epidemiology of chronic pancreatitis: burden of the disease and consequences. United Eur Gastroenterol J. 2014; 2:345–54.

4. Éva Kereszturi, Richárd Szmola, Zoltán Kukor et al. Hereditary pancreatitis caused by mutation induced misfolding of human cationic trypsinogen - a novel disease mechanism. Hum Mutat. 2009; 30(4): 575–582.

5. Zhang Y-F, Deng H-L, Fu J, Zhang Y, Wei J-Q. Pancreatitis in hand-foot-And-mouth disease caused by enterovirus 71. World J Gastroenterol. 2016; 22(6):2149–52.

6. Sarner M, Cotton PB. Classification of pancreatitis. Gut. 1984; 25:756–9.

7. Barman KK, Premalatha G, Mohan V. Tropical chronic pancreatitis. Postgrad Med J. 2003; 79:606–15

8. Paliwal S, Bhaskar S, Chandak GR. Genetic and phenotypic heterogeneity in tropical calcific pancreatitis. World J Gastroenterol. 2014; 20(46):17314–23.

9. Zahid Hassan, Viswananthan Mohan, Liaquat Ali et al. SPINK1 Is a Susceptibility Gene for Fibrocalculous Pancreatic Diabetes in Subjects from the Indian Subcontinent. Am J Hum Genet. 2002; 71(4): 964–968

10. Olson OC, Joyce JA. Cysteine cathepsin proteases: regulators of cancer progression and therapeutic response. Nature Reviews Cancer. 2015; 15(12):712–29.

11. Pungerčar JR, Caglič D, Sajid M, Dolinar M, Vasiljeva O, Požgan U, et al. Autocatalytic processing of *procathepsin B* is triggered by proenzyme activity. FEBS journal. 2009; 276(3):660–8.

12. Katunuma N. Posttranslational processing and modification of cathepsins and cystatins. J Signal Transduct. 2010; 375–345.

13. Ghosh P, Dahms NM, Kornfeld S. Mannose 6-phosphate receptors: new twists in the tale. Nature reviews Molecular cell biology. 2003; 4(3):202–13.

14. Pungercar JR, Caglic D, Sajid M, Dolinar M, Vasiljeva O, Pozgan U, et al. Autocatalytic processing of *procathepsin B* is triggered by proenzyme activity. FEBS J. 2009; 276(3):660–8.

15. Lerch MM, Halangk W. Human pancreatitis and the role of *cathepsin B*. Gut. 2006; 55(9):1228–30.

16. Gasteiger E, Hoogland C, Gattiker A, Duvaud Se, Wilkins MR, Appel RD, et al. Protein identification and analysis tools on the ExPASy server: Springer. 2005.

17. Mahurkar S, Idris MM, Reddy D, Bhaskar S, Rao G, Thomas V, et al. Association of *cathepsin B* gene polymorphisms with tropical calcific pancreatitis. Gut. 2006; 55(9):1270–5.

18. Šali A, Blundell TL. Comparative protein modelling by satisfaction of spatial restraints. Journal of molecular biology. 1993;234(3):779–815.

19. Eisenberg D, Lüthy R, Bowie JU. VERIFY3D: assessment of protein models with threedimensional profiles. Methods in enzymology. 1997; 277:396.

20. Wiederstein M, Sippl MJ. ProSA-web: interactive web service for the recognition of errors in three-dimensional structures of proteins. Nucleic acids research 35(suppl 2). 2007; W407–W10.

21. S.C. Lovell, I.W. Davis, W.B. Arendall III, P.I.W. de Bakker, J.M. Word, M.G. Prisant, J.S. Richardson and D.C. Richardson. Structure validation by Calpha geometry: phi,psi and Cbeta deviation. Proteins: Structure, Function & Genetics. 2002; 50: 437–450.

22. Zhang Y, Skolnick J. TM-align: a protein structure alignment algorithm based on the TM-score. Nucleic acids research. 2005; 33(7):2302–9.

23. Ng PC, Henikoff S. SIFT: Predicting amino acid changes that affect protein function. Nucleic acids research. 2003; 31(13):3812–4.

24. Adzhubei IA, Schmidt S, Peshkin L, Ramensky VE, Gerasimova A, Bork P, et al. A method and server for predicting damaging missense mutations. Nature methods. 2010; 7(4):248–9.

25. Mi H, Muruganujan A, Casagrande JT, Thomas PD. Large-scale gene function analysis with the PANTHER classification system. Nature protocols. 2013; 8(8):1551–66.

26. Willard L, Ranjan A, Zhang H, Monzavi H, Boyko RF, Sykes BD, et al. VADAR: a web server for quantitative evaluation of protein structure quality. Nucleic acids research. 2003; 31(13):3316–9.

27. Worth CL, Preissner R, Blundell TL. SDM—a server for predicting effects of mutations on protein stability and malfunction. Nucleic acids research. 2011.

28. Capriotti E, Fariselli P, Casadio R. I-Mutant2.0: predicting stability changes upon mutation from the protein sequence or structure. Nucleic acids research. 2005; 33 (Web Server issue): W306–W310.

29. Douglas E. V. Pires, David B. Ascher and Tom L. Blundell. mCSM: predicting the effects of mutations in proteins using graph-based signatures. Bioinformatics. 2014; 30(3): 335–342

30. H.J.C. Berendsen, D. van der Spoel, R. van Drunen. GROMACS: A message-passing parallel molecular dynamics implementation. Computer Physics Communications. 1995; 91:43–56.

31. Andrew E. Bruno, Li Li, James L. Kalabus, Yuzhuo Pan, Aiming Yu, Zihua Hu miRdSNP: a database of disease-associated SNPs and microRNA target sites on 3’UTRs of human genes. BMC Genomics. 2012; 13(1):44.

32. Bhattacharya A, Ziebarth JD, Cui Y. PolymiRTS Database 3.0: linking polymorphisms in microRNAs and their target sites with human diseases and biological pathways. Nucleic Acids Res. 2014; 42(D1):D86–D91.

33. Jing Gong, Yin Tong, Hong-Mei Zhang, An-Yuan Guo. miRNASNP: a database of miRNA related SNPs and their effects on miRNA function. BMC Bioinformatics. 2012; 13(Suppl 18):A2

34. Flavio Mignone, Carmela Gissi, Sabino Liuni and Graziano Pesole. Untranslated regions of mRNAs. Genome Biology. 2002; 3(3).

35. Podobnik M, Kuhelj R, Turk V, Turk D. Crystal structure of the wild-type human *procathepsin B* at 2.5 Å resolution reveals the native active site of a papain-like cysteine protease zymogen. Journal of molecular biology. 1997; 271(5):774–88.

36. Orlichenko L, Stolz DB, Noel P, Behari J, Liu S, Singh VP. ADP-ribosylation factor 1 protein regulates trypsinogen activation via organellar trafficking of *procathepsin B* protein and autophagic maturation in acute pancreatitis. Journal of Biological Chemistry. 2012; 287(29):24284–93.

37. Penner RC, Andersen ES, Jensen JL, Kantcheva AK, Bublitz M, Nissen P, Rasmussen AM, Svane KL, Hammer B, Rezazadegan R, Nielsen NC. Hydrogen bond rotations as a uniform structural tool for analyzing protein architecture. Nature communications. 2014; 17;5

